# Yeast-expressed Recombinant SARS-CoV-2 Receptor Binding Domain, RBD203-N1 as a COVID-19 Protein Vaccine Candidate

**DOI:** 10.1101/2021.08.24.457518

**Authors:** Wen-Hsiang Chen, Jeroen Pollet, Ulrich Strych, Jungsoon Lee, Zhuyun Liu, Rakhi Tyagi Kundu, Leroy Versteeg, Maria Jose Villar, Rakesh Adhikari, Junfei Wei, Cristina Poveda, Brian Keegan, Aaron Oakley Bailey, Yi-Lin Chen, Portia M. Gillespie, Jason T. Kimata, Bin Zhan, Peter J. Hotez, Maria Elena Bottazzi

**Author notes:** correspondence: Maria Elena Bottazzi, 1102 Bates St., Ste. 550 | Houston, TX 77030.

## Abstract

**Background:** SARS-CoV-2 protein subunit vaccines are being evaluated by multiple manufacturers to fill the need for low-cost, easy to scale, safe, and effective COVID-19 vaccines for global access. Vaccine candidates relying on the receptor-binding domain (RBD) of the SARS-CoV-2 spike protein have been the focus of our development program. In this paper, we report on the generation of the RBD203-N1 yeast expression construct, which produces a recombinant protein that when formulated with alum and the TLR-9 agonist, CpG1826 elicits a robust immune response and protection in mice.

**Method:** The RBD203-N1 antigen was expressed in the yeast *Pichia pastoris* X33. After fermentation at the 5 L scale, the protein was purified by hydrophobic interaction chromatography followed by anion exchange chromatography. The purified protein was characterized biophysically and biochemically, and after its formulation, the immunogenicity and efficacy were evaluated in mice.

**Results, Conclusions, and Significance:** The RBD203-N1 production process yielded 492.9 ± 3.0 mg/L of protein in the fermentation supernatant. A two-step purification process produced a >96% pure protein with a recovery rate of 55 ± 3% (total yield of purified protein: 270.5 ± 13.2 mg/L fermentation supernatant). The protein was characterized as a homogeneous monomer with well-defined secondary structure, thermally stable, antigenic, and when adjuvanted on alum and CpG, it was immunogenic and induced robust levels of neutralizing antibodies against SARS-CoV-2 pseudovirus. These characteristics show that this vaccine candidate is well suited for technology transfer with feasibility of its transition into the clinic to evaluate its immunogenicity and safety in humans.

## 1. INTRODUCTION

As of August 20^th^, 2021, close to 5 billion doses of coronavirus vaccines have been administered in over 180 countries. However, this impressive vaccination campaign has still left over 70% of the global population without access to efficient protection from COVID-19 [1]. According to a recent analysis, people in the highest-income countries are getting vaccinated more than 20 times faster than those living in poverty [2]. Therefore, there remains an urgent need to add additional safe and effective vaccines to the global inventory and to produce these vaccines at the lowest cost possible when it comes to production, storage, and distribution.

Recombinant protein expression in yeast is a low-cost and therefore attractive platform of production as compared to other more costly production systems for biologics such as mammalian cell culture systems [3]. This has been demonstrated for multiple vaccine antigens in general [4, 5], and is currently the case for additional COVID-19 vaccines under development. The Argentinian AntiCovid Consortium, for example, showed recently that a SARS-CoV-2 receptor-binding domain antigen was just as well folded and stable when made in yeast as when it was produced in mammalian cell culture [6]. Another yeast-produced RBD when displayed on hepatitis B virus-like particles was shown to effectively reduce viral loads in the respiratory tract of immunized cynomolgus macaques [7].

Our group has previously shown that a yeast-produced RBD vaccine antigen candidate (amino acid residues 331-549 of the SARS-CoV-2 spike protein), when combined with alum and 3M-052 (TLR7/8 agonist), was able to protect *Rhesus macaques* from challenge with SARS-CoV-2 by eliciting robust humoral and cellular immune responses [8]. To reduce hyperglycosylation, aggregation, improve stability and enable better controlled scalable and reproducible process development, we removed one of the main glycosylation sites (N331) from the RBD and mutated a C-terminal cysteine residue (C538A). The resulting protein, RBD219-N1C1, was shown to maintain its ability to effectively trigger a robust immune response with a high level of neutralizing antibodies against SARS-CoV-2 [9, 10].

Here we report on the design, construction, and biophysical, biochemical, and immunological evaluation of a new construct, RBD203-N1 (residues 332–l533), where we deleted the SARS CoV-2 RBD residues 534-549, including the cysteine residue at position 538. The data reported here support the potential of an RBD203-N1 protein-based vaccine as a candidate for technology transfer and its suitability for its transition into the clinic to evaluate safety, immunogenicity, and efficacy in humans.

## 2. MATERIALS AND METHODS

### 2.1. Cloning and Fermentation of SARS-CoV-2 RBD203-N1 in *Pichia pastoris*

The recombinant *Pichia pastoris* X-33 construct expressing RBD203-N1 (residues 332–533 of the SARS-CoV-2 spike protein, GenBank: QHD43416.1) was generated as described previously [9, 11]. In short, the DNA encoding RBD203-N1 was synthesized and subcloned into the *Pichia* secretory expression vector pPICZαA (Invitrogen) using EcoRI/XbaI restriction sites (GenScript). The recombinant plasmid was transformed into *P. pastoris* X-33.

The RBD203-N1 pPICZαA/P. *pastoris X33* construct was fermented in 5 L vessels [9, 11, 12]. Briefly, the glycerol seed stock was used to inoculate 0.5 L Buffered Minimal Glycerol (BMG) medium for overnight culture, which was then used to inoculate 2.5 L sterile low salt medium (LS) in a fermenter containing 3.5 mL/L PTM1 trace elements and 3.5 mL/L 0.02% d-Biotin. Fermentation was initiated at 30 °C and pH 5.0, with dissolved oxygen (DO) maintained at 30%. Upon DO spike, the pH was ramped up to 6.5 using 14% ammonium hydroxide, and the temperature was lowered to 25°C over 1 hour. Induction was initiated by adding methanol from 1 mĻ/L/h to 11 mĻ/L/h over 6 hours. After the methanol adaption stage, induction was maintained at 25 °C with a methanol feed rate from 11 to 15 for another 64 hours [12]. After fermentation, the culture was harvested by centrifugation. The fermentation supernatant (FS) was filtered using a 0.45 μm PES filter and evaluated by SDS-PAGE.

### 2.2. Protein Purification

RBD203-N1 was purified based on Process-2 described in Lee *et al.* [12] with slight modifications in the capture step. Ammonium sulfate was added to the FS to a final concentration of 1.1 M (w/v) followed by pH adjustment to 8.0, and filtration through a 0.45 μm PES filter. The filtered material was loaded onto a 51.5 mL Butyl Sepharose HP column (Cytiva), which was washed with buffer A (30 mM Tris-HCl pH 8.0) containing 1.1 M ammonium sulfate. Bound protein was eluted in buffer A containing 0.44 M ammonium sulfate. UFDF and a polish step followed as described in the original *Process-2* [12]. Protein yield and the purity for the in-process and final purified RBD203-N1 were analyzed by SDS-PAGE. As a protein control, the yeast expressed RBD219-N1C1 protein was used and generated in-house as described [12].

### 2.3. Western Blot

Two micrograms of RBD203-N1 or RBD219-N1C1 were loaded on 4-20% Tris-glycine gels, and transferred to a polyvinylidene difluoride membrane, and probed with eight different in-house generated mouse monoclonal antibodies raised against SARS-CoV-2 RBD219-WT (1μg/mL in 10mL; mAB #s 1128, 643, 486, 902, 854, 942, 748 and 102), respectively. A 1:3,000 dilution of an AP-conjugated goat anti-mouse IgG (KPL) was used as the secondary antibody.

### 2.4. ELISA using Anti-RBD2l9-N1C1 Mouse Monoclonal Antibodies

In this experiment, we evaluated the binding of eight anti-SARS-CoV-2 RBD219-WT mAbs (# 1128, 643, 486, 902, 854, 942, 748, and 102) to RBD203-N1 and RBD219-N1C1. Ninety-six-well ELISA plates were coated with 100 μL 2 μg/mL of either RBD203-N1 or RBD219-N1C1 overnight in duplicate at 4°C followed by blocking with PBST/0.1% BSA overnight at 4°C. Once the plates were blocked, 100 μL 3x serially-diluted mAb with an initial concentration of 2 μg/mL was added to the wells. The plates were incubated at room temperature for 2 hours to allow mAb to bind to RBDs. After this binding step, the plates were washed with PBST four times followed by adding 100 μL 1:6,000 diluted HRP conjugated anti-mouse IgG antibodies (LSBiosciences) and incubated for 1 hour at room temperature. Finally, 100 μL TMB substrate was added and incubated for 4 minutes in the dark to react with HRP. The reaction was terminated with 100 μL of 1M HCl and absorption readings were taken at 450 nm using a BioTek EPOCH 2 microplate reader.

### 2.5. Identity and purity by SE-HPLC

Waters Alliance HPLC Separations Modules and Associated PDA Detectors were operated as per the vendor’s instruction. Fifty micrograms of Bio-Rad gel filtration standard or RBD203-N1 were injected into a TSKgel® G2000SWXL column (300 mm X 7.8 mm), and eluted in 20 mM Tris, 150 mM NaC1, pH 7.5 (1X TBS) at the flow rate of 0.6 mL/min.

### 2.6. Size Assessment by Dynamic Light Scattering (DLS)

The size of RBD203-N1 in solution was analyzed using DLS. Briefly, the concentration of the protein was adjusted to 1 mg/mL using 1X TBS. The samples were then filtered through 0.02 μm filters. Four replicates of forty microliters of protein were loaded into each well of a clear bottom 384-well plate. The hydrodynamic radii of the proteins were measured using a DynaPro Plate Reader II.

### 2.7. Structural Assessment by Circular Dichroism (CD)

Purified RBDs were diluted with deionized water to a final concentration of 0.2 mg/mL and loaded into a 0.1 cm path cuvette. Dilution with water was to reduce the chloride ion content, which is known to interfere with the CD absorbance, especially at low wavelengths. CD spectra were obtained from 250 to 190 nm with a Jasco J-1500 spectrophotometer set at 100 nm/min and a response time of 1 s at 25°C. The CD data were analyzed using a CD Analysis and Plotting Tool (https://capito.uni-jena.de/index.php). In addition, the RBDs (0.5 mg/mL) were heated from 25 °C to 95 °C for a denaturation profile analysis.

### 2.8. Structural Assessment by Thermal shift

RBD203-N1 or RBD219-N1C1 were diluted to 0.32 mg/mL and mixed with the reagents in Protein Thermal Shift™ Dye kit (Thermo Fisher) as per the vendor’s instructions. In short, 12.5 μL of 0.32 mg/mL RBD were mixed with 5 μL of Protein Thermal Shift buffer, followed by 2.5 μL of 8x Protein Thermal Shift dye in three to four replicates. These samples were vortexed briefly and centrifuged to remove any bubbles and further heated from 25 °C to 95 °C to monitor the change of fluorescence intensity using a ViiA™ 7 Real-Time PCR system.

### 2.9. *in vitro* Functionality Assay by ELISA (ACE-2 binding)

Ninety-six-well ELISA plates were coated with 100 μL 5 μg/mL ACE-2-hFc (LakePharma) overnight at 4°C followed by blocking with PBST/0.1% BSA. Once the plates were blocked, 100 μL serially diluted RBD219-N1C1 or RBD203-N1 with an initial concentration of 40 μg/mL were added to the wells. The plates were incubated at room temperature for 2 hours to allow ACE-2 to interact with each RBD. After this binding step, the plates were washed with PBST four times followed by adding 100 μL of 1:5,000 diluted anti-RBD219-WT horse sera followed by 1:10,000 diluted HRP conjugated anti-horse IgG antibodies and incubating for 1 hour at room temperature. Finally, 100 μL TMB substrate were added and incubated for 15 min in the dark to react with HRP. The reaction was terminated with 100 μL of 1M HCl and absorption readings were taken at 450 nm using a BioTek EPOCH 2 microplate reader.

### 2.10. Preclinical Study Design

A preclinical study in mice was performed under the approved Institutional Animal Care and Use Committee (IACUC) protocol at Baylor College of Medicine. The study design is shown in Supplementary Table 1. Formulations were prepared with 7 μg protein per dose, and the protein was first adsorbed on 200 μg of aluminum hydroxide (alum; containing 100 μg of aluminum) before 20 μg of CpG1826 (vac-1826-1, Invivogen) were added at the point of injection. 6–8-week-old Female BALB/c mice were immunized twice intramuscularly (i.m.) at 21-day intervals and then euthanized 14 days after the second immunization.

### 2.11. Antigen-specific Antibody Measurements by ELISA

To examine RBD-specific antibodies in mouse sera, indirect ELISAs were conducted as described previously [13]. Briefly, 96-well ELISA plates were coated with 100 μL of 2 μg/mL RBDs in 1x coating buffer and incubated overnight at 4 °C. The plates were then blocked with 200 μL/well PBST/0.1% BSA for 2 hours at room temperature. After being washed once with 300 μL PBST. 100 μL of serially diluted mouse serum samples, naïve mouse serum, and blank (PBST/0.1% BSA) were added to the plate and incubated for 2 hours at room temperature. The plates were further washed four times with PBST and dispensed with 100 μL of 1:6,000 diluted goat anti-mouse IgG HRP for 1 hour at room temperature, followed by washing five times with PBST. Finally, 100 μL TMB substrate were added to each well and incubated for 15 minutes at room temperature. After incubation, the reaction was stopped by adding 100 μL of 1 M HCl. The absorbance at a wavelength of 450 nm was measured using a BioTek Epoch 2 spectrophotometer.

### 2.12. Cytokine Measurements by Luminex

Splenocytes preparation and cytokine measurements were performed as previously described [13]. Briefly, GentleMACS Octo Dissociator was used to dissociate spleen and pelleted splenocytes. The splenocytes were then resuspended in 1 mL ACK lysing buffer for 1 minute at room temperature followed by the addition of 40 mL PBS. Splenocytes were again pelleted and resuspended in 5 mL 4°C cRPMI (RPMI 1640 + 10% HI FBS + 1x pen/strep) and transferred through a 40 μm filter to obtain a single-cell suspension.

For the in vitro cytokine release assay, splenocytes were seeded in a 96-well culture plate at 1×10^6^ live cells in 250 μL cRPMI and stimulated with 10 μg/mL RBDs for 48 hours at 37° C 5% CO_2_, PMA/Ionomycin and media were used as the positive and negative control, respectively. After incubation, 96-well plates were centrifuged and the supernatant was transferred to a new 96-well plate to measure levels of IL-1β, IL-2, IL-4, IL-6, IL-10, IL-13, IL-17A, IFN-γ, and TNF-α using Milliplex Mouse Th17 Luminex kit (EMD Millipore) on a MagPix Luminex instrument. Raw data were first analyzed by Bio-Plex Manager software followed by Excel and Prism.

### 2.13. Pseudovirus assay

Pseudovirus experiments were executed as previously published [13]. Using in vitro grown human 293T-hACE2 cells, infected cells were quantified based on the expression of luciferase. The plasmids used for the pseudovirus production are the luciferase-encoding reporter plasmid (pNL4-3.lucR-E-), Gag/Pol-encoding packaging construct (pΔ8.9), and codon-optimized SARS-CoV-2 spike protein expression plasmids (pcDNA3.1-CoV-2 S gene) based on clone p278-1. Pseudovirus containing supernatants were recovered after 48 hours and passed through a 0.45 μm filter and saved at −80°C until used for neutralization studies.

Ten microliters of pseudovirus (~500 relative infection units) were incubated with serial dilutions of the serum samples for 1 hour at 37°C. Next, 100 μL of sera-pseudovirus were added to 293T-hACE2 cells in 96-well poly-D-lysine coated culture plates. Following 48 hours of incubation in a 5% CO2 environment at 37°C, the cells were lysed with 100 μL of Promega Glo Lysis buffer for 15 min at RT. Finally, 50 μL of the lysate were added to 50 μL luc substrate (Promega Luciferase Assay System). The amount of luciferase was quantified by luminescence (relative luminescence units (RLU)), using the Luminometer (Biosynergy H4). The percentage (%) virus inhibition was calculated as

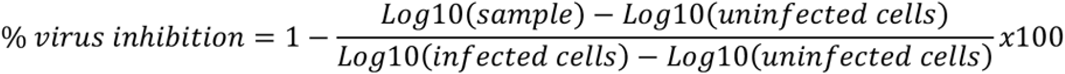

Serum from vaccinated mice was also compared by their 50% inhibitory dilution (IC50), defined as the serum dilution at which the virus infection was reduced by 50% compared with the negative control (virus + cells).

## 3. RESULTS

### 3.1. Cloning, Production, Size and Purity Evaluation of RBD203-N1

SARS-CoV-2 RBD203-N1 (residues 332-533 of the spike protein) is a truncated version of the previously developed SARS-CoV-2 vaccine antigen, RBD219-N1C1, with 16 amino acid residues removed from the C-terminus. N1 designates the exclusion of N331, a putative N-glycosylation site, from the construct (**Figure 1**). To evaluate the reproducibility of the production process, two identical 5L scale production runs were performed. During production, the yield and the recovery were monitored (**Table 1**). The results indicated a fermentation yield for RBD203-N1 of 492.9 ± 3.0 mg/L of fermentation supernatant (FS) with an overall recovery of 55 ± 3% after purification. When evaluating the coefficient of variation of the process, one could notice that the %CV was lower than 6% throughout the process, indicating that the process was reproducible. Purity analysis of the in-process samples (**Figure 2A**) revealed that the downstream process efficiently improved the purity from 61.8 ± 1.1% to 97.0 ± 0.4% under reduced conditions, or from 75.0 ± 0.6% to 96.4 ± 0.9% under non-reduced conditions. SE-HPLC data also revealed the purity of RBD203-N1 was approximately 99.9% (**Figure 2B**). Additionally, DLS indicated that RBD203-N1 was monodispersed (5.9% Polydispersity) with an estimated molecular weight of 31 kDa (**Figure 2C**).

**Figure 1.**
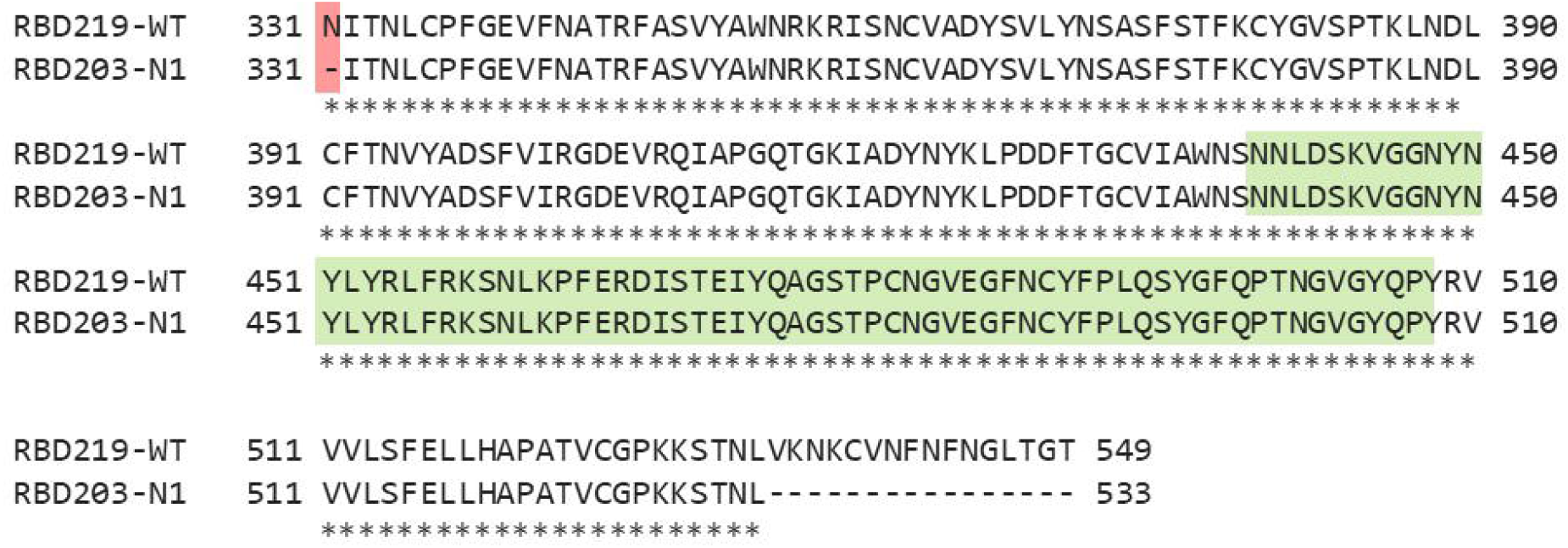
Sequence alignment between RBD219-WT, and RBD203-N1 of the SARS-CoV-2 spike protein. N1 designated the exclusion of N331 (highlighted in red). The region highlighted in green is the receptor-binding motif.

**Table 1.** Purification Yield and Process Recovery for RBD203-N1

| Step | Yield (mg/L of FS)<sup>a</sup> |  |  |  | Step Recovery (%)<sup>a</sup> |  |  |  | Overall Recovery (%)<sup>a</sup> |  |  |  |
| --- | --- | --- | --- | --- | --- | --- | --- | --- | --- | --- | --- | --- |
|  | Run 1 | Run 2 | Average | %CV | Run 1 | Run 2 | Average | %CV | Run 1 | Run 2 | Average | %CV |
| FS | 495.0 | 490.7 | 492.9±3.0 | 0.6 | - | - | - | - | 100 | 100 | 100±0 | 0.0 |
| Capture<br>(Butyl HP) | 327.1 | 322.7 | 324.9±3.1 | 1.0 | 66 | 66 | 66±0 | 0.0 | 66 | 66 | 66±0 | 0.0 |
| UFDF<br>(Pellicon<br>XL) | 323.9 | 329.6 | 326.8±4.0 | 1.2 | 99 | 102 | 101±2 | 2.0 | 65 | 67 | 66±1 | 1.5 |
| Polish<br>(QXL) | 261.2 | 279.8 | 270.5±13.2 | 4.9 | 81 | 85 | 83±3 | 3.6 | 53 | 57 | 55±3 | 5.5 |

<sup>a</sup> Average ± SD from two independent purification runs

**Figure 2.**
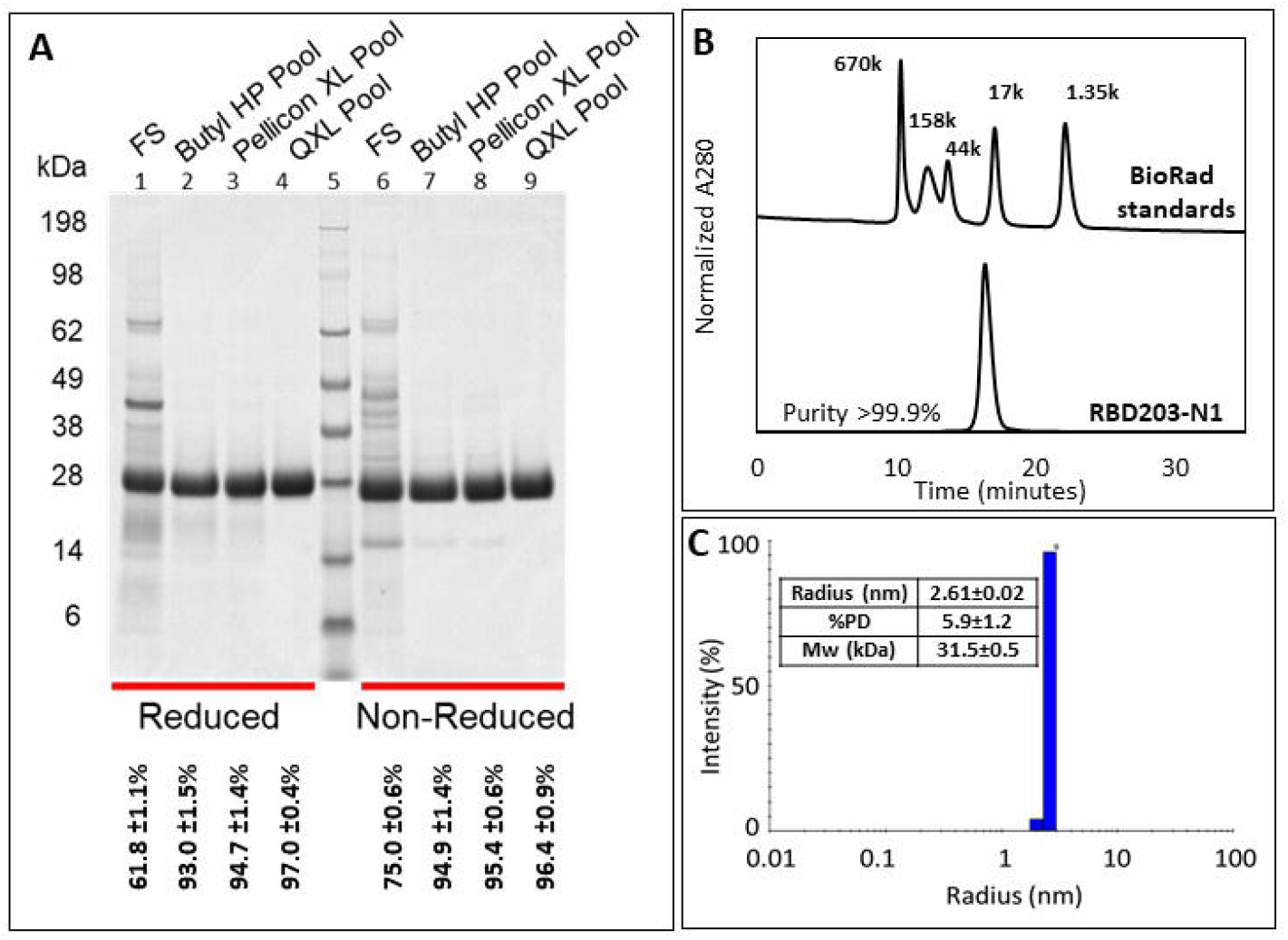
(A) Purity assessment of in-process samples by SDS-PAGE (A), Purity and size assessment of purified RBD203-N1 by SE-HPLC (B) and by DLS (C).

### 3.2. Western Blot and ELISA with monoclonal antibodies

The antigenicity of RBD203-N1 against 8 different in-house generated anti-RBD monoclonal antibodies were evaluated using western blot with RBD219-N1C1 as a control (**Figure 3**). Overall, the binding profile of the antibodies to RBD203-N1 and RBD219-N1C1 was similar. Neutralizing antibodies (mAbs 1128, 643, and 486) likely recognized conformational epitopes and thus did not recognize reduced RBDs well. mAbs 854 and 942 recognized both non-reduced and reduced RBD equally, while mAbs 748 and 102 recognized the reduced RBDs stronger than the reduced RBDs. Although 203 dimer was not detectable in SDS-PAGE, SE-HPLC, these monoclonal antibodies all recognized the RBD203 dimer form. Interestingly, mAb486 only recognized the RBD dimer but not the monomer, suggesting that the dimer form might have better preserved the conformation for antibody recognition.

**Figure 3.**
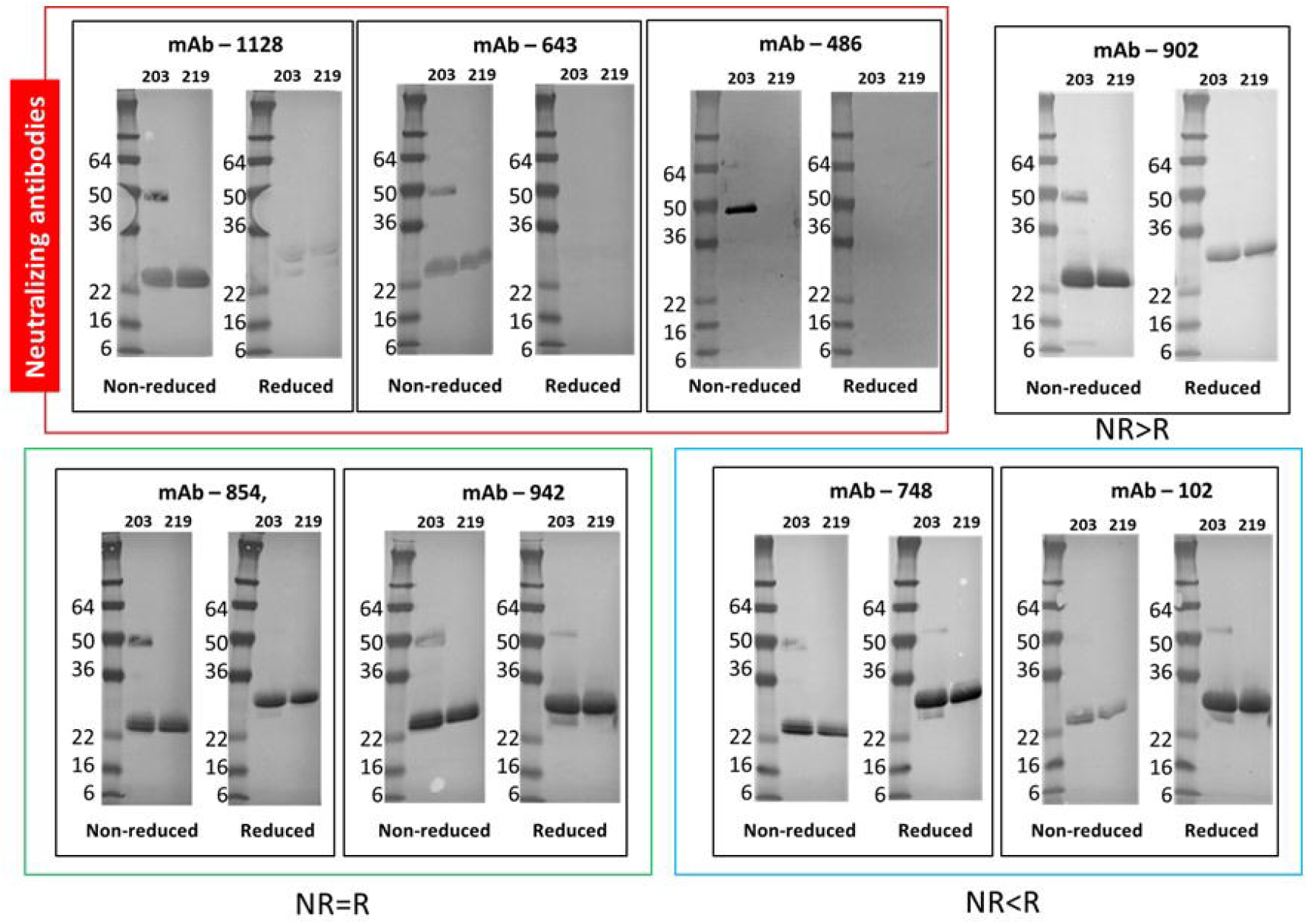
Western blot analysis for RBD203-N1 (203) and RBD219-N1C1 (219) using eight anti-RBD219-N1C1 Mouse Monoclonal Antibodies.

Similar to the western blot, ELISAs were performed using the same monoclonal panel against RBD203-N1 and RBD219-N1C1, respectively. Similar binding profiles were observed for both proteins for seven of the eight mAbs in a native condition. With mAb-486, a slightly lower affinity to RBD203-N1 was observed (**Figure 4**).

**Figure 4.**
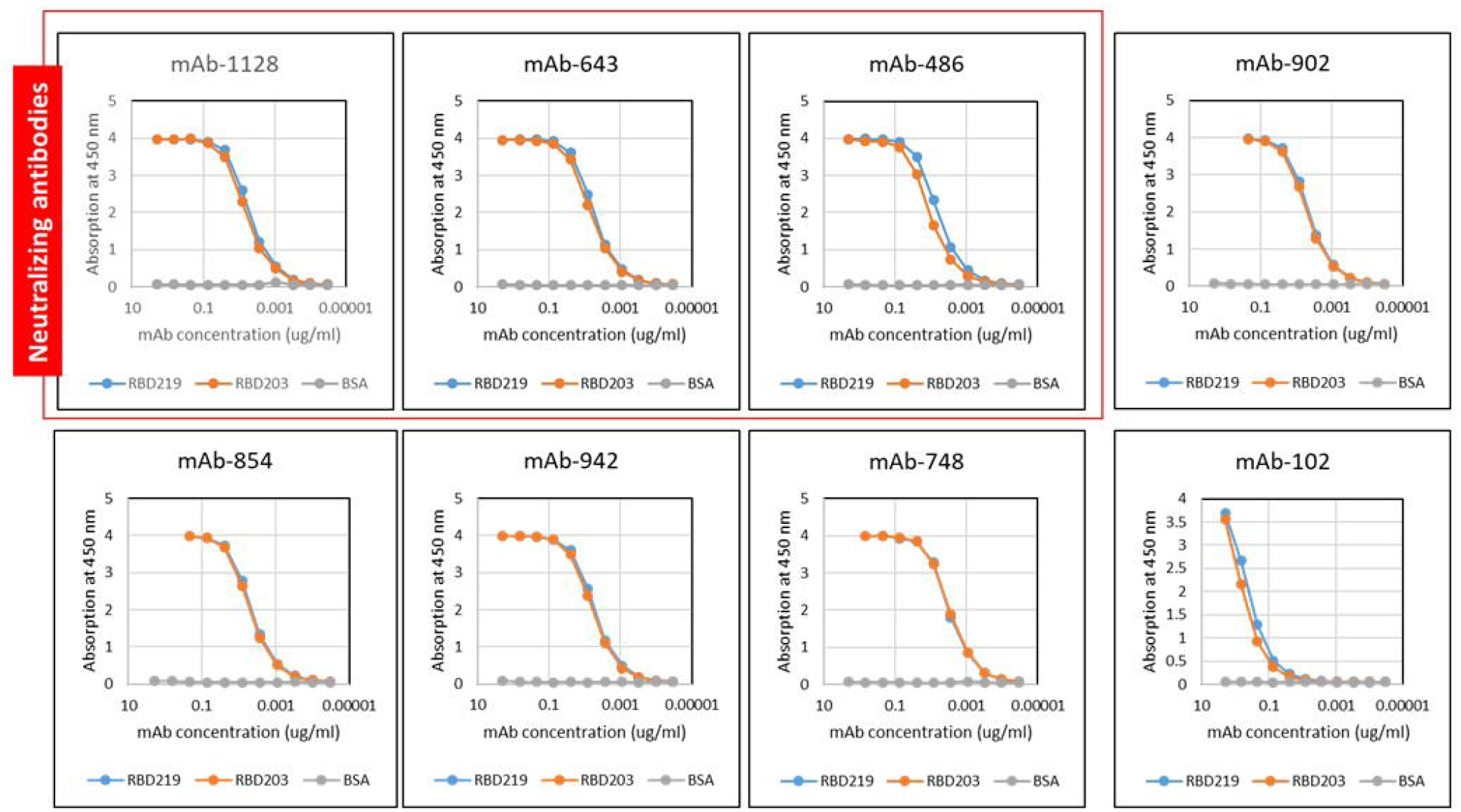
Monoclonal antibody ELISA for RBD203-N1 (RBD203) and RBD219-N1C1 (RBD219). BSA was used as a negative control. RBDs or BSA were first coated on the plate, followed by incubation with specific RBD-mAbs, as indicated on top of each panel. Binding of the mAbs was detected using an HRP-labeled anti-mouse IgG secondary mAb.

### 3.3. Secondary structure thermal stability assessment

When far-UV CD spectrometry was performed to investigate the secondary structure of RBD203-N1 in comparison with RBD219-N1C1, we observed very similar data (**Figure 5A**). The thermal stability of the secondary structures was evaluated by heating the samples from 25 °C to 95 °C (**Figures 5B-5C**) and CD melting curves and their derivatives were further examined at 231 nm (**Figures. 5D-5E**). Based on the derivative, the average melting temperatures (Tm) were 50.8 °C and 51.9 °C for RBD203-N1 and RBD219-N1C1, respectively, suggesting similar thermal stability.

**Figure 5.**
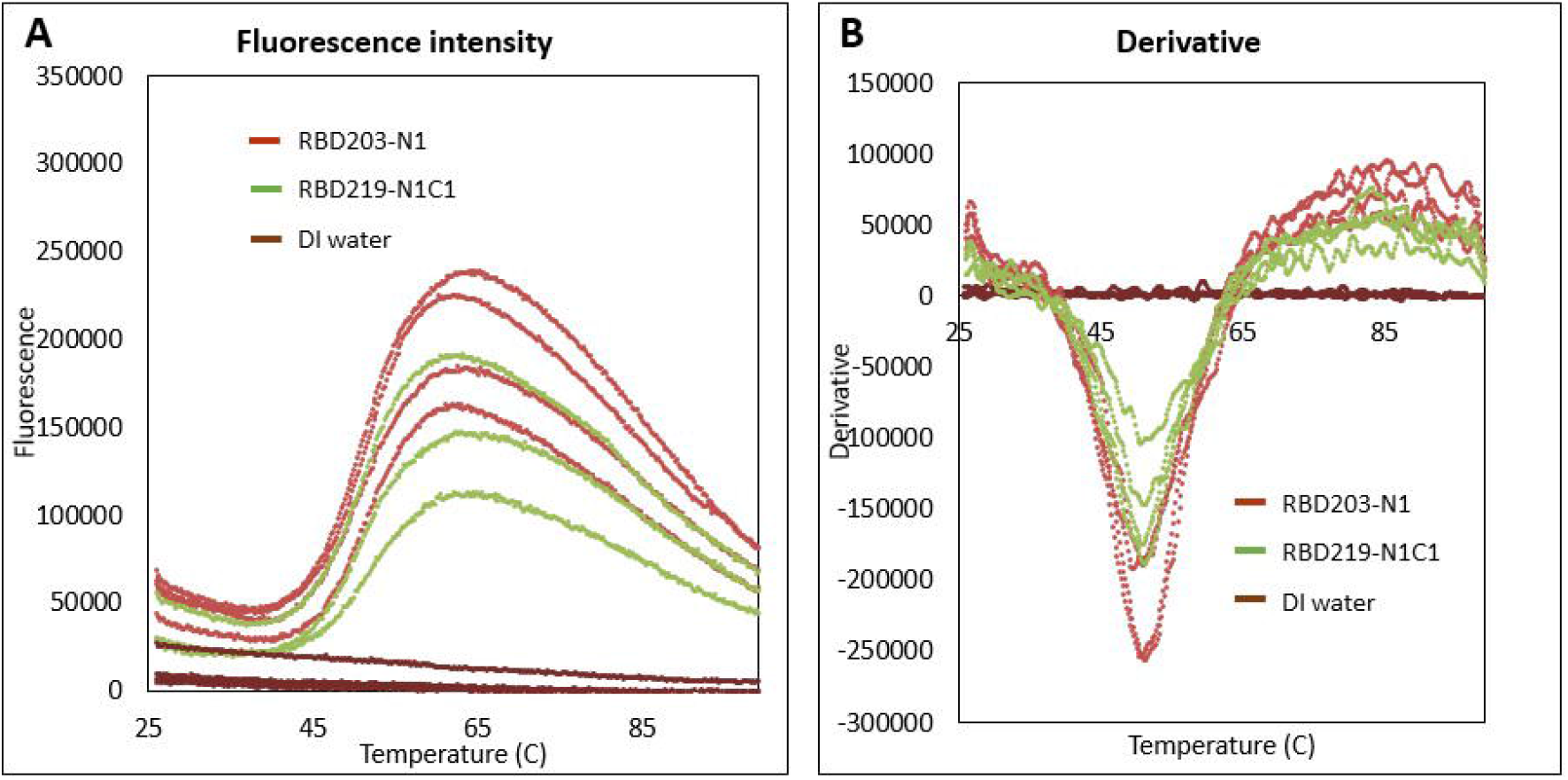
Circular dichroism (CD) analysis of RBD203-N1 and RBD219-N1C1, including CD profile (A), overall melting profile of RBD203-N1 (B), and RBD219-N1C1 (C), and CD readouts and derivatives of RBD203-N1 (D) and RBD219-N1C1 (E) at 231 nm extracted from the overall melting profile.

### 3.4. Tertiary structure thermal stability assessment

In this study, we used thermal shift assays to compare the thermal stability of the tertiary structure for RBD203-N1 and RBD219-N1C1. The melting curve (**Figure 6A**) showed a similar fluorescence profile between these two RBDs. The initial fluorescence of both proteins indicated similar surface hydrophobicity when they were still intact. When the temperature was increased, these two proteins started to denature (T_on_) at approximately 38 °C. Calculated from the derivatives, the melting temperatures (Tm) were 50.4. ± 0.6 °C and 50.7 ± 0.2 °C for RBD203-N1 and RBD219-N1C1, respectively (**Figure 6B**), which further suggested that these two RBDs shared similar tertiary structures.

**Figure 6.**
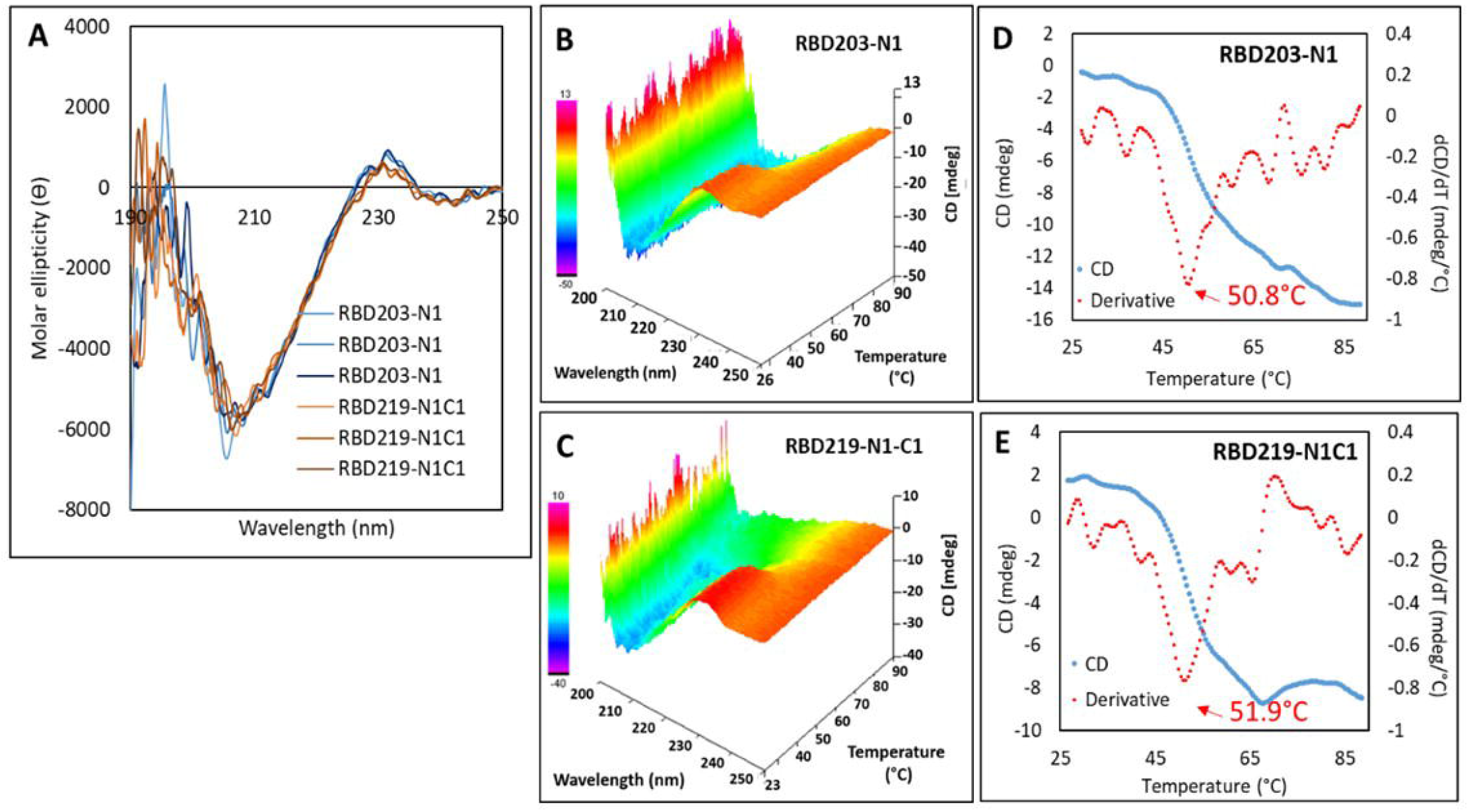
Thermal shift assay to investigate the tertiary structure for RBD203-N1 and RBD219-N1C1 (A) and the derivative fluorescence-temperature plot (B). Water was used as a negative control.

### 3.5. The RBD203-N1 protein efficiently binds to ACE-2 *in vitro*

Li *et al.* have indicated that the most potent neutralizing antibodies that recognized the RBD blocked its binding to ACE-2 [14] and thus, confirming the ability of RBD to bind to ACE-2 is crucial. When comparing RBD203-N1 and RBD219-N1C1 in this way, both RBDs bound to ACE-2 similarly with EC50 values of 0.0417 ± 0.005 μg/mL and 0.0410 ± 0.004 μg/mL, respectively (**Figure 7**).

**Figure 7.**
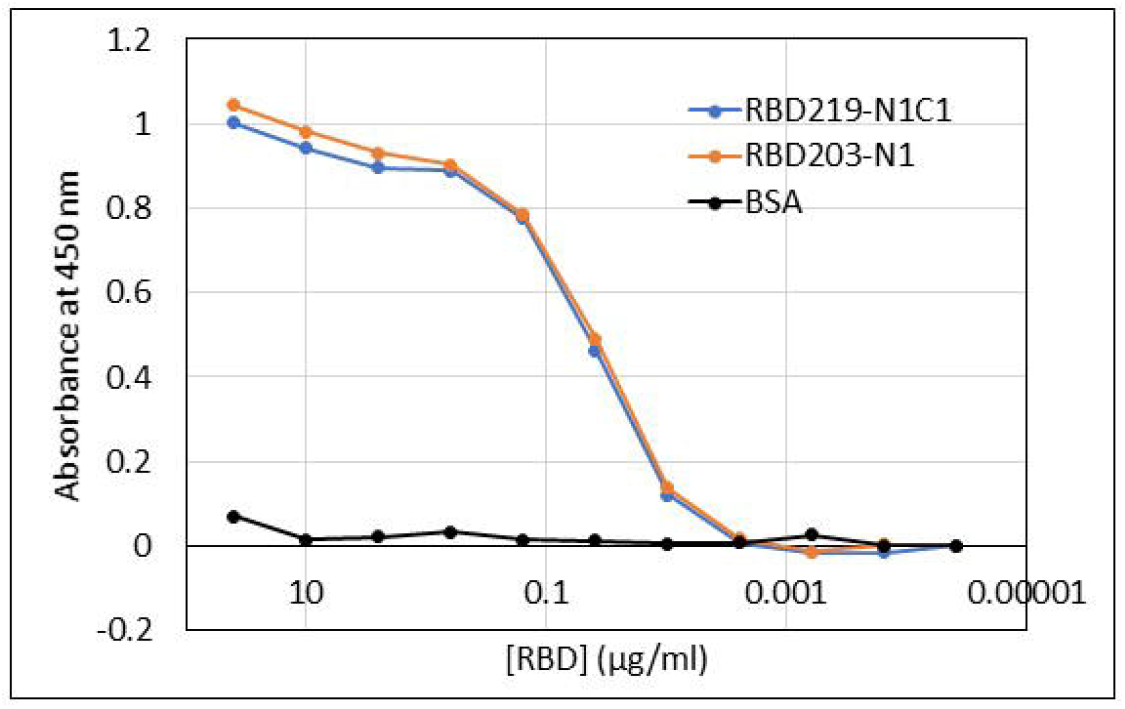
ACE-2 binding study to evaluate the functionality of RBD203-N1 and RBD219-N1C1

### 3.6. RBD203-N1 formulated with alum/CpG triggered strong immunity and neutralizing activity

The study design to evaluate the immunogenicity and neutralizing activity is shown in **Figure 8A**. Mice were vaccinated twice on days 0 and 21, and on day 35, serum was tested for total anti-RBD IgG (**Figure 8B**). With the addition of 20 μg of CpG in the formulation, we observed an approximately 1,000-fold increase in the magnitude of the total IgG titer and a noticeable reduction of the intra-cohort variation for both proteins. Luminex assays were used to evaluate the levels of cytokines after restimulation of splenocytes with RBD N1C1, and the heatmap (**Figure 8C**) indicated both RBD203-N1 and RBD219-N1C1 triggered similar cytokine profiles with the same formulations. Consistent with the data previously shown [10], when formulated with alum alone, secretion of IFN-gamma, IL-6, and IL-10 was observed, while the addition of CpG produced a stronger and more balanced Th1/Th2 response, with increased levels of IL-2, IL-4, IL-6 and IFN-gamma(**Figure 8C**).

**Figure 8:**
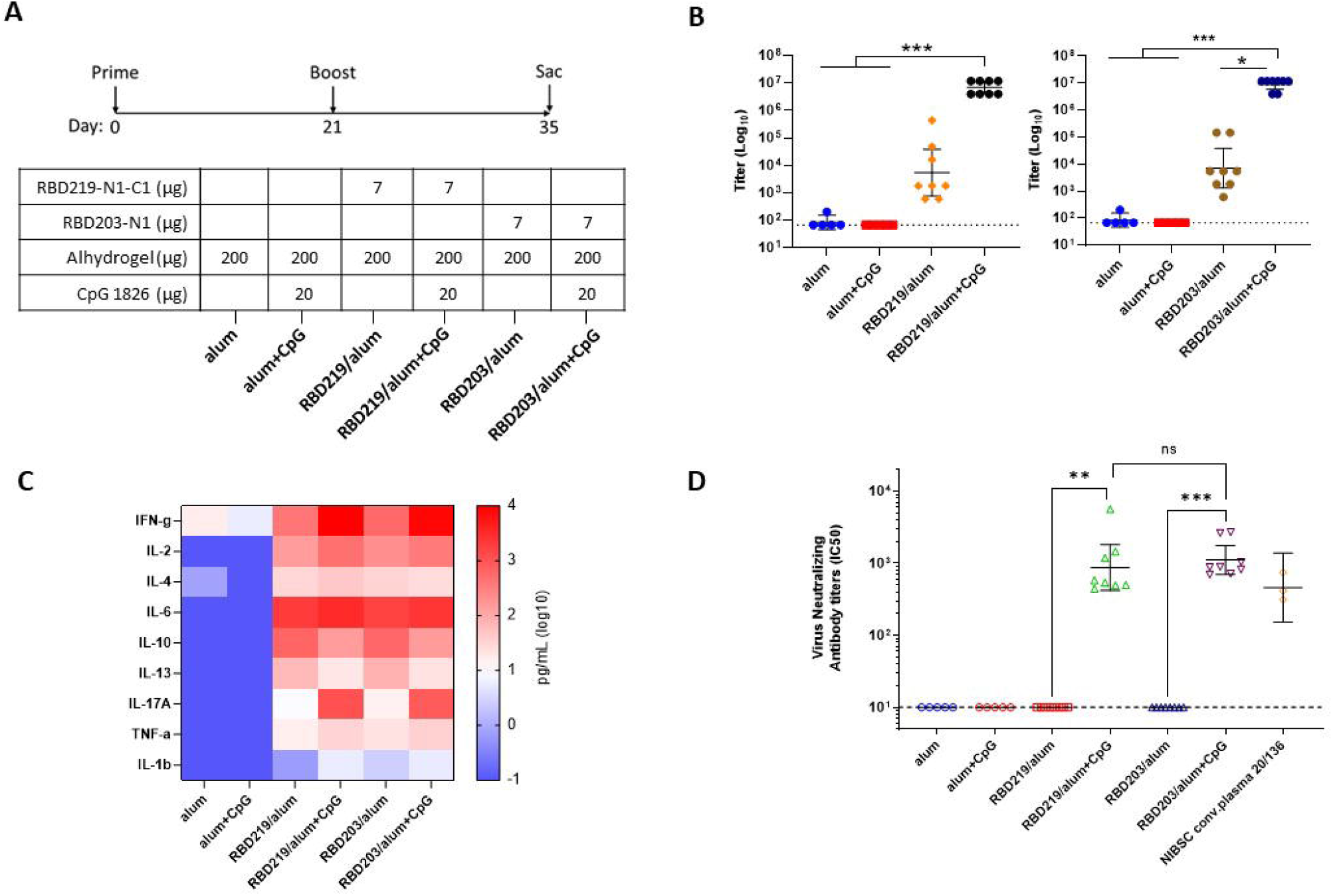
A) Study timeline and table with vaccine formulations, B) Total IgG titers measured from mouse sera against RBD219-N1C1 protein (left) or RBD203-N1 protein (right). C) Heatmap of secreted cytokines measured in supernatant from splenocytes re-stimulated with RBD219-N1C1. Median values were calculated within each treatment group for each cytokine. D) IC50 values were measured by a neutralization assay using a lentiviral SARS-CoV-2 Wuhan pseudovirus. Kruskal Wallis tests were performed to evaluate for statistical significance between different groups. p> 0.12 (not significant, ns), p < 0.033 (*),p < 0.002 (**), p<0.001 (***). RBD203 stands for RBD203-N1 and RBD219 stands for RBD219-N1C1.

When neutralizing capacity was evaluated in a pseudovirus assay (**Figure 8D**), no neutralizing antibodies were detected in the sera of mice immunized twice with RBD/alum, alum, and alum+CpG. However, mice immunized with two doses of the 7 μg RBD/alum+CpG showed approximately 2.5-fold higher neutralizing antibody titers than the National Institute for Biological Standards and Control (NIBSC) human convalescent plasma standard. No significant differences in the titers were observed between mice immunized with RBD203-N1 or RBD219-N1C1, formulated alum+CpG. Collectively, the data suggest that RBD203-N1 and RBD219-N1C1 elicit similar levels of antigen-specific antibodies, neutralizing antibodies, and cytokines.

## 4. DISCUSSION

Here we report on a COVID-19 vaccine candidate antigen based on a truncated receptor-binding domain construct of the SARS-CoV-2 spike protein. The antigen, RBD203-N1, was expressed effectively in the yeast *P. pastoris*, and purified by a combination of hydrophobic interaction chromatography and anion exchange chromatography. The fermentation yield using a process that we have previous developed for a similar vaccine antigen [12], was determined as 492.9 ± 3.0 mg/L RBD203-N1 of FS. The overall recovery from this two-step purification scheme for RBD203-N1 was determined as 55 ± 3%. The production process was demonstrated to be reproducible with less than 6% of %CV throughout the process between two identical production runs. RBD203-N1 was shown to be a protein of high purity when analyzed by SDS-PAGE (>96%) and SE-HPLC (>99%). DLS also indicated that the purified protein was monodispersed. When we inspected the molecular weight of the deglycosylated RBD203-N1 by mass spectrometry (**Supplementary method and Supplementary Figure 1**), we discovered two major RBD203-N1 species with additional EAEAEF or EAEF amino acid residues at the N-terminus. The EAEA residues are expected remnants due to well-described inefficient *P. pastoris* Ste13-protease cleavage of the signal peptide upstream of recombinant proteins expressed in the pPicZα/P. *pastoris* system [15]. The adjacent EF residues are derived from the translation of the EcoRI used for cloning of the RBD sequence in pPiczα. Nevertheless, the different purified RBD203-N1 lots were consistently 82-85% of the EAEAEF-variant and 15-18% of the EAEF-variant again proving the reproducibility of the production process.

We further assessed the biophysical and biochemical characteristics of RBD203-N1 by evaluating its antigenicity, secondary structure, thermal stability, and *in vitro* functionality. When eight in-house monoclonal antibodies, generated against the wild-type RBD219 (no deletion of N or C) [8, 9] were used to evaluate the antigenicity of these two RBDs by western blot and ELISA, both RBDs were recognized mostly to the same extent. One exception was mAb486 that only recognized the dimer form of RBD203-N1 in the western blot, which suggested that the dimer form might have preserved the confirmation better. When assessing the secondary structures, far-UV CD spectra indicated that RBD203-N1 and RBD219-N1C1 proteins had similar secondary structures and the melting temperatures evaluated by CD further revealed that the thermal stability for the secondary structure of both proteins was comparable. Additionally, thermal shift assays also indicated that both RBDs shared comparable thermal stability for their tertiary structures. Moreover, the in vitro functionality assay further confirmed a similar binding affinity to ACE-2 to these RBDs, suggesting that these two RBDs shared the same biophysical and biochemical characteristics.

The immunogenicity in mice of RBD203-N1, when formulated with alum with or without the TLR9 agonist CpG, was evaluated. The addition of CpG to COVID-19 vaccine formulations has been demonstrated to promote antigen dose sparing as well as the induction of balanced Th1/Th2 immune responses with much lower intra-cohort variability [13]. When adjuvanted with CpG, the use of 7ug and 2.3ug of RBD203-N1 protein elicited robust neutralizing antibody titers that were protective against SARS-CoV-2 pseudovirus particles. The level of neutralizing antibodies in the serum was 2.5-times higher than the NIBSC standard plasma and was also equivalent to the control RBD219-N1C1/alum+CpG vaccine [13]. The use of low RBD protein concentration, when formulated with alum alone and in a two-dose regime, did not trigger robust antigen-specific antibody titers and the neutralizing activity was undetectable. However, our studies with RBD/alum formulations in two- or three-dose regimens using higher protein doses have been shown to trigger robust immune responses with high neutralizing titers [10]. In addition, Nanogen, recently showed that their Nanocovax vaccine, consisting of a recombinant S protein formulated with alum was immunogenic and efficacious in various animal models [16]. Therefore, RBD proteins, including RBD203-N1, adjuvanted with alum alone should continue to be evaluated for safety, immunogenicity, and efficacy especially in the context of the changing SARS CoV-2 virus epidemiology.

## 5. Conclusions

In this study, we report on RBD203-N1, a truncated version of the SARS CoV-2 spike protein RBD. The fermentation yield of this construct was 493 mg/L of FS. The two-step purification process allowed for a recovery of more than 50% of RBD203-N1. The purified RBD203-N1 was of high purity (>96% by SDS-PAGE and >99% by SE-HPLC). When studying the biophysical and biochemical characteristics, we confirmed this truncated protein retained the expected secondary structure, thermal stability, antigenicity, and functionality. Additionally, when formulated with alum+CpG, it triggered a robust level of antigen-specific antibodies that possess neutralizing ability, as well as a desired balanced cytokine profile. Collectively, the data suggested that RBD203-N1 is a suitable vaccine candidate antigen for technology transfer and transition into the clinic to evaluate its safety, immunogenicity, and efficacy in humans.

## ABBREVIATIONS

COVID-19: Coronavirus disease 2019
SARS: severe acute respiratory syndrome
CoV: coronavirus
S: spike
RBD: receptor-binding domain
DO: dissolved oxygen
FS: fermentation supernatant
CV: column volume
%CV: coefficient of variation
DLS: dynamic light scattering
CD: circular dichroism
ACE-2: angiotensin-converting enzyme 2
AEX: anion exchange chromatography
HIC: hydrophobic interaction chromatography
i.m.: intramuscular
SE-HPLC: size-exclusion high-performance liquid chromatography
CpG: CpG oligodeoxynucleotide adjuvant
mAB: monoclonal antibody
TLR: toll-like receptor
NIBSC: National Institute for Biological Standards and Control

## Author contributions

WHC and JBP conceived the study, designed and performed experiments, interpreted data and drafted the manuscript; US designed experiments, interpreted data and drafted the manuscript; JL, ZL, LV, BZ designed and performed experiments, interpreted data, and reviewed the manuscript; RTK, MJV, RA, JW, CP, BK, AOB, YLC, BL performed experiments and reviewed the manuscript; PMG and JTK designed experiments and reviewed the manuscript; PJH and MEB conceived the study, designed experiments, provided scientific guidance and reviewed the manuscript.

## Acknowledgments

This work was supported by the Robert J. Kleberg Jr. and Helen C. Kleberg Foundation; Fifth Generation, Inc. (Tito’s Handmade Vodka); JPB Foundation, NIH-NIAID (AI14087201); and Texas Children’s Hospital Center for Vaccine Development Intramural Funds. We also would like to thank PATH Center for Vaccine Innovation and Access (Seattle, WA, USA) for their guidance as well as technical and intellectual support. The Mass Spectrometry Facility at the University of Texas Medical Branch is supported in part by Cancer Prevention Research Institute of Texas (CPRIT), grant number RP190682.

