## Supplementary for "Yeast-expressed Recombinant SARS-CoV-2 Receptor Binding Domain, RBD203-N1 as a COVID-19 Protein Vaccine Candidate"

### SUPPLEMENTARY DATA

#### Supplementary Method

#### Intact Mass Analysis of Deglycosylated Proteins

RBD203-N1 or RBD219-N1C1 were treated with PNGase-F (NEB) following the vendor's manual to remove N-linked glycans. The deglycosylated proteins were then analyzed by micro-flow reversed-phase LC-MS using a 45-min gradient with an Ultimate 3000 UHPLC system coupled to Orbitrap Fusion mass spectrometer (ThermoFisher Scientific), operated at a resolution setting of 240,000 (at m/z 200). The data were analyzed using the Xtract deconvolution algorithm in Biopharma Finder v4.0 (ThermoFisher Scientific).

**Supplementary Table 1**. Study group and formulation information. N1C1: RBD219-N1C1; 203-N1: RBD2023-N1.

| Group Label | Group Size | Mouse # | Antigen | Antigen Dose (µg) | Adjuvant Molecule | Adjuvant Dose (µg) | Route | Volume (mL) |
|---|---|---|---|---|---|---|---|---|
| **1** | 5 | 1-5 | None | - | Alhydrogel® | 200 | IM | 0.05 |
| **2** | 5 | 6-10 | None | - | Alhydrogel®+CpG | 200 + 20 | IM | 0.05 |
| **3** | 8 | 11-18 | N1C1 | 25 | Alhydrogel® | 200 | IM | 0.05 |
| **4** | 8 | 19-26 | 203-N1 | 25 | Alhydrogel® | 200 | IM | 0.05 |
| **5** | 8 | 27-34 | N1C1 | 7 | Alhydrogel® | 200 | IM | 0.05 |
| **6** | 8 | 35-42 | N1C1 | 7 | Alhydrogel®+CpG | 200 + 20 | IM | 0.05 |
| **7** | 8 | 43-50 | 203-N1 | 7 | Alhydrogel® | 200 | IM | 0.05 |

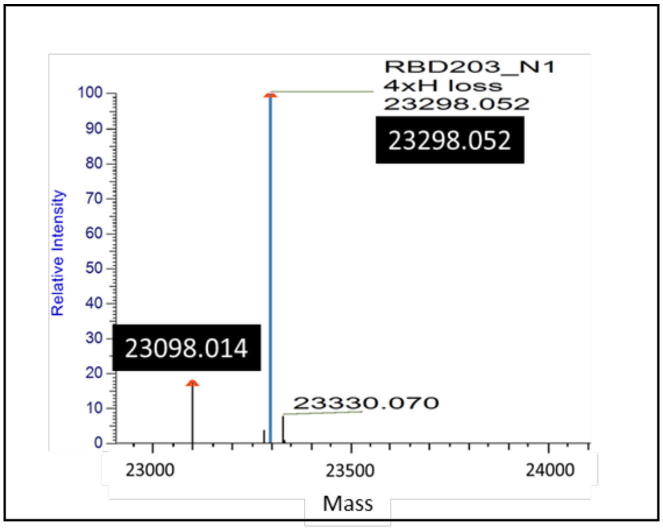


**Supplementary Figure 1.** Intact mass spectrometry for deglycosylated RBD203-N1. This result indicated two major species matching the sequences of (1) RBD203-N1[a] with additional EAEF amino acid residues at the N-terminus (23,098 Da), and (2) RBD203-N1[b] with additional EAEAEF amino acid residues at the N-terminus (23,298 Da). 82-85% of the RBD203-N1 is the EAEAEF variant while 15-18% is the EAEF variant.
